# HiC-Spector: A matrix library for spectral and reproducibility analysis of Hi-C contact maps

**DOI:** 10.1101/088922

**Authors:** Koon-Kiu Yan, Galip Guürkan Yardimci, William S. Noble, Mark Gerstein

## Abstract

**Summary:** Genome-wide proximity ligation based assays like Hi-C have opened a window to the 3D organization of the genome. In so doing, they present data structures that are different from conventional 1D signal tracks. To exploit the 2D nature of Hi-C contact maps, matrix techniques like spectral analysis are particularly useful. Here, we present HiC-spector, a collection of matrix-related functions for analyzing Hi-C contact maps. In particular, we introduce a novel reproducibility metric for quantifying the similarity between contact maps based on spectral decomposition. The metric successfully separates contact maps mapped from Hi-C data coming from biological replicates, pseudo-replicates and different cell types.

**Availability:** Source code in Julia and the documentation of HiC-spector can be freely obtained at https://github.com/gersteinlab/HiC_spector

## 1 Introduction

Genome-wide proximity ligation assays such as Hi-C have emerged as powerful techniques to understand the 3D organization of the genome (Lieberman-Aiden et al., 2009; Kalhor et al., 2011). While these techniques offer new biological insights, they demand different data structures and present new computational questions (Dekker et al., 2013; Ay and Noble, 2015). For instance, a basic question of particular practical importance is, how can we quantify the similarity between two Hi-C data sets? In particular, given two experimental replicates, how can we determine if the experiments are really reproducible?

Data from Hi-C experiments are usually summarized by so-called chromosomal contact maps. By binning the genome into equally sized bins, a contact map is a matrix whose elements store the population-averaged co-location frequencies between pairs of loci. Therefore, mathematical tools like spectral analysis can be extremely useful in understanding these chromosomal contact maps. The aim of this project is to provide a set of basic analysis tools for handling Hi-C contact maps. In particular, we introduce a simple but novel metric to quantify the reproducibility of the maps using spectral decomposition.

## 2 Algorithms

We represent a chromosomal contact map by a symmetric and non-negative adjacency matrix *W*. The matrix elements represent the frequencies of contact between genomic loci and therefore serve as a proxy of spatial distance. The larger the value of *W_ij_*, the closer is the distance between loci *i* and *j*. The starting point of spectral analysis is the Laplacian matrix *L*, which is defined as *L = D-W*. Here *D*is a diagonal matrix in which *D_ii_ = Σ_i_ W_ij_* (the coverage of bin *i* in the context of Hi-C). As in many other applications, the Laplacian matrix further takes a normalized form *ℒ = D^-1/2^LD^-1/2^* (Chung, 1997). It can be verified that 0 is an eigenvalue of *ℒ*, and the set of eigenvalues of *ℒ* (0<λ_0_<λ_1_<…<λ_n-1_) is referred to as the spectrum of *ℒ*. Like common dimensionality reduction procedures, the first few eigenvalues are of particular importance because they capture the basic structure of the matrix, whereas the very high eigenvalues are essentially noise. The normalized Laplacian matrix is closely related to random walk processes taking place in the underlying graph of *W*. In fact, the first few eigenvalues correspond to the slower decay modes of the random walk process, and capture the large-scale structure of the contact map.

Given two contact maps *W*^A^ and *W*^B^, we propose to quantify their similarity by decomposing their corresponding Laplacian matrices *ℒ*^A^ and *ℒ*^B^ respectively and then comparing their eigenvectors. Let 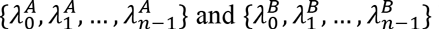 be the spectra of *ℒ*^A^ and *ℒ*^B^, and 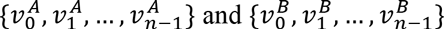 be their sets of normalized eigenvectors. A distance metric *S*_d_ is defined as

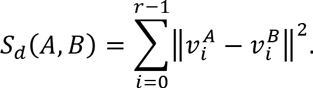

Here ║.║ represents the Euclidean distance between the two vectors. The parameter r is the number of leading eigenvectors picked from *ℒ^A^* and *ℒ^B^*. In general, *S*_d_ provides a metric to gauge the similarity between two contact maps. 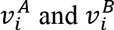 are more correlated if A and B are two biological replicates as compared to the case when they are two different cell lines (see Supplemental Information).

For the choice of r, like any principal component analysis, the contribution of leading eigenvectors is more important than the contribution from lower ranked eigenvectors. In fact, we observe that the Euclidean distance between a pair of high-order eigenvectors is the same as the distance between a pair of unit vectors whose components are randomly sampled from a standard normal distribution (see Supplemental Information). In other words, the high-order eigenvectors are essentially noise terms, whereas the signal is stored in the leading vectors. As a rule of thumb, we found the choice *r* = 20 is good enough for practical purposes. Furthermore, as the distance between a pair of randomly sampled unit vectors presents a natural limit 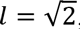, we rescale the distance metric into a reproducibility score Q ranges from 0 to 1 by

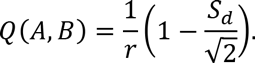

As shown in Figure 1, the reproducibility scores between pseudo-replicates are greater than the scores for real biological replicates, which are greater than the scores between maps from different cell lines.

**Figure 1.**
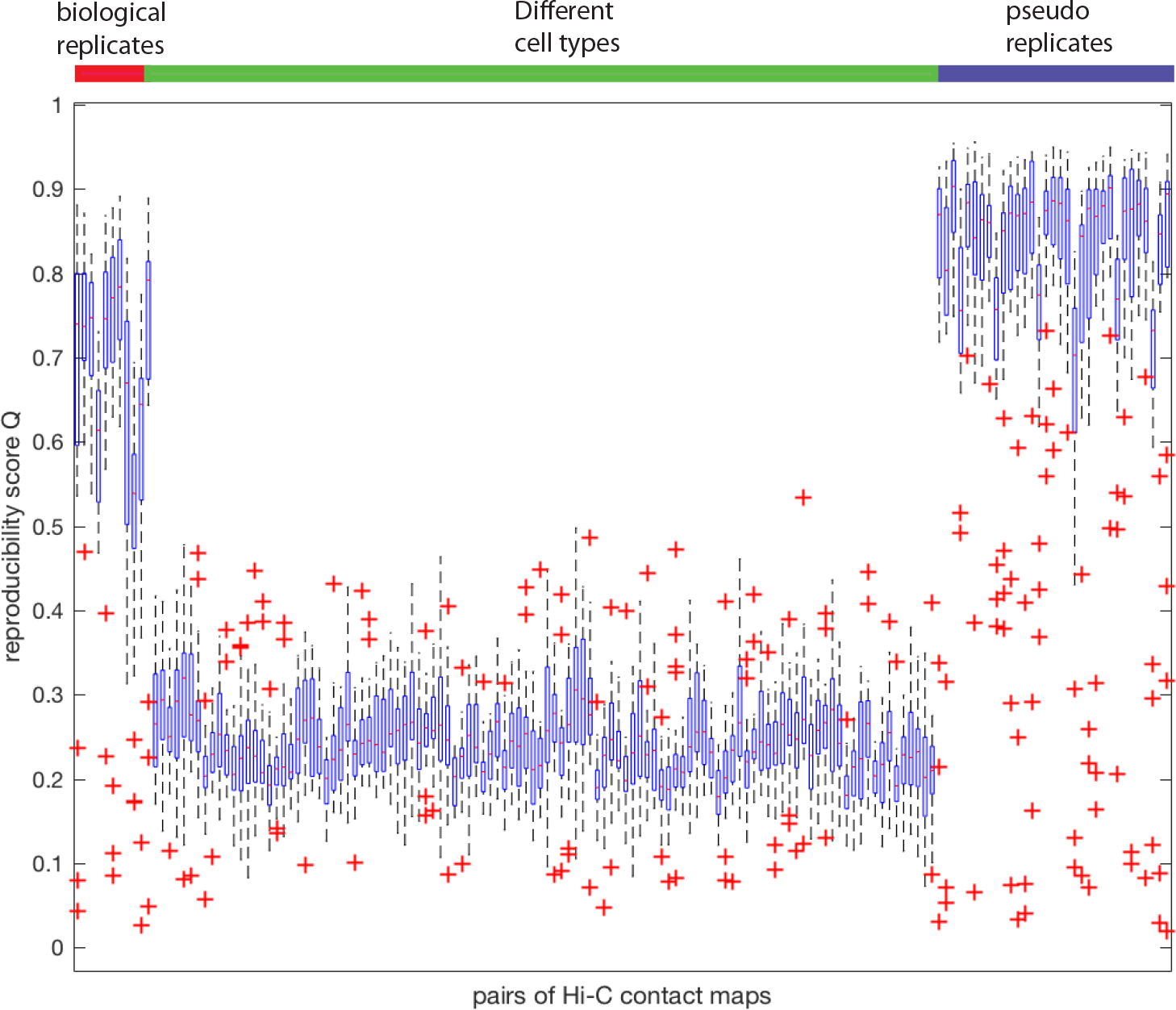
Reproducibility scores for 3 sets of Hi-C contact maps pairs. Contact maps came from Hi-C experiments performed in 11 cancer cell lines by the ENCODE consortium (https://www.encodeproject.org/). Biological replicates refer to a pair of replicates of the same experiment. Pseudo replicates are obtained by pooling the reads from two replicates together and performing down sampling. There are 11 biological replicates, 33 pairs of pseudo replicates, and 110 pairs of maps between different cell types. Each box shows the distribution of Q in 23 chromosomes.

Apart from the reproducibility score, a number of matrix algorithms useful for analyzing contact maps are provided in HiC-spector. For instance, we have a function for performing matrix balancing using the Knight-Ruiz algorithm (Knight and Ruiz, 2012), which is widely used as a normalization procedure for contact maps (Imakaev et al., 2012). In addition, we have included the functions for estimating the average contact frequency with respect to the genomic distance, as well as identifying the so-called A/B compartments (Lieberman-Aiden et al., 2009) using the corresponding correlation matrix.

## 3 Implementation and Benchmark

HiC-spector is a library written in Julia, a high-performance language for technical computing. The bottleneck for evaluating the reproducibility metric we introduced is matrix diagonalization. The runtime depends very much on the size of contact maps. We found the performance efficient for most practical purposes, for instance, given a pair of contact maps of human chromosome 1 with bin-size equal to 40kb, it takes 80 seconds on a laptop with 2.8GHz Intel Core i7 and 16Gb of RAM.

## Acknowledgements

We want to thank the 3D Nucleome subgroup in the ENCODE consortium for processing the Hi-C data and discussion.

## Supporting Information

**Figure S1:**
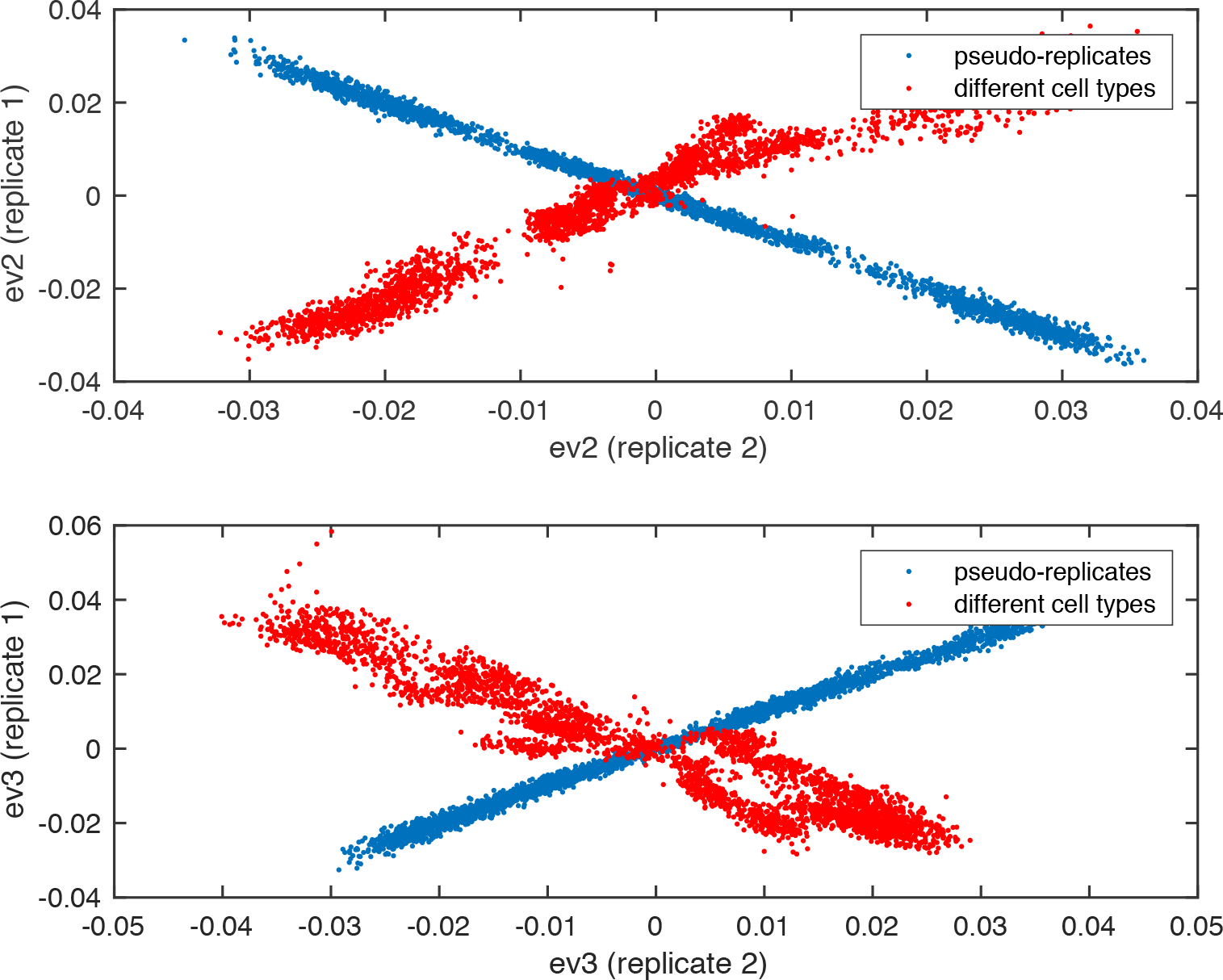
Leading eigenvectors of contact maps. Blue refers to a pair of pseudo-replicates. The corresponding leading eigenvectors are more correlated as compared to red, which refers to a pair of contact maps originating from two different cell lines.

**Figure S2:**
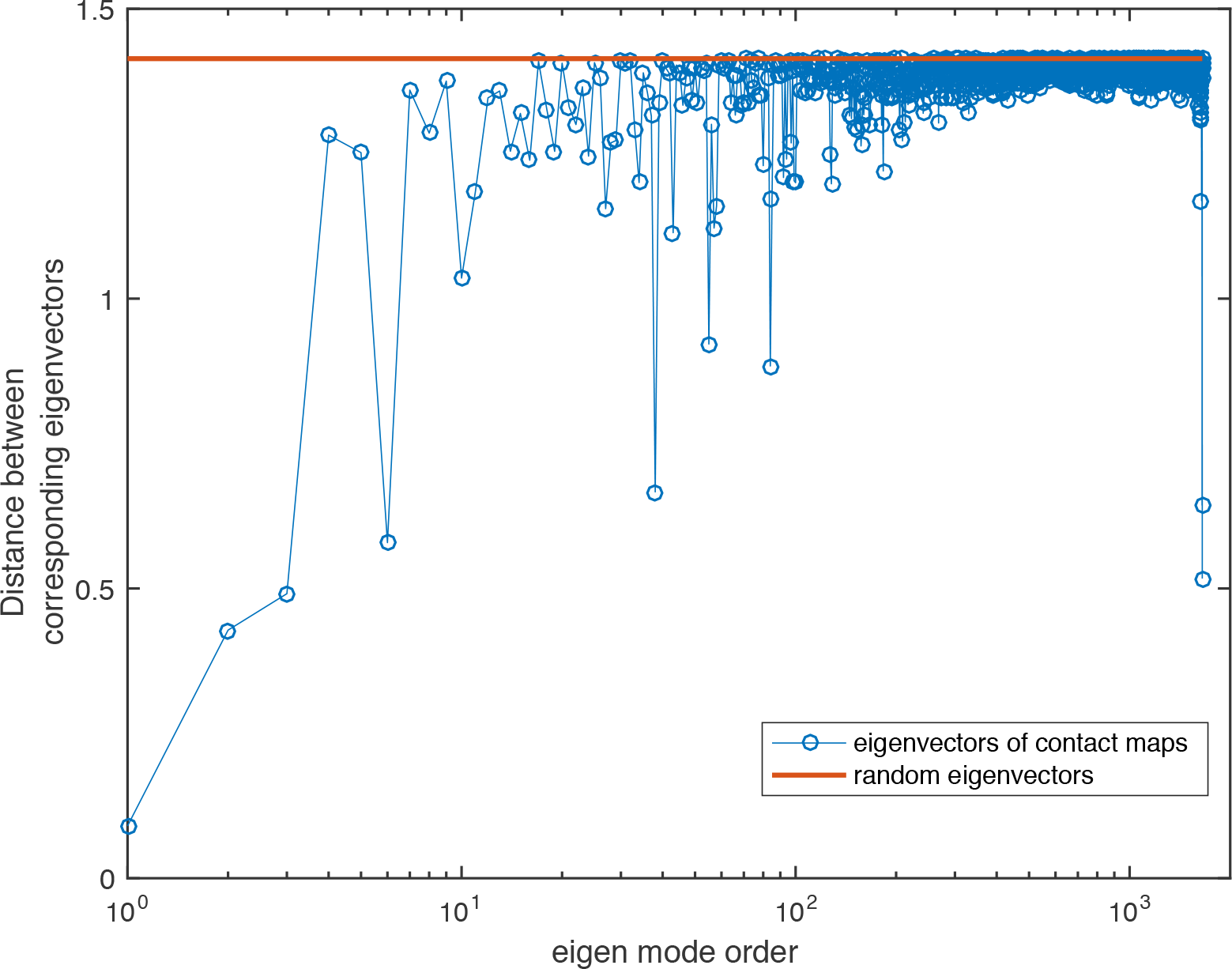
Euclidean distance between corresponding eigenvectors in a pair of Hi-C contact maps. The distance between leading eigenvectors is low. The red line is the distance between two random unit vectors whose components are sampled from a standard normal and then rescaled. The distance between two high-order eigenvectors is very close to the red line, suggesting they are noise instead of actual signal.

## References

Ay, F., and Noble, W.S. (2015). Analysis methods for studying the 3D architecture of the genome. Genome Biol. 16, 183.

Chung, F. (1997). Spectral graph theory (American Mathematical Society).

Dekker, J., Marti-Renom, M.A., and Mirny, L.A. (2013). Exploring the three-dimensional organization of genomes: interpreting chromatin interaction data. Nat. Rev. Genet. 14, 390–403.

Imakaev, M., Fudenberg, G., McCord, R.P., Naumova, N., Goloborodko, A., Lajoie, B.R., Dekker, J., and Mirny, L.A. (2012). Iterative correction of Hi-C data reveals hallmarks of chromosome organization. Nat. Methods 9, 999–1003.

Kalhor, R., Tjong, H., Jayathilaka, N., Alber, F., and Chen, L. (2011). Genome architectures revealed by tethered chromosome conformation capture and population-based modeling. Nat. Biotechnol. 30, 90–98.

Knight, P.A., and Ruiz, D. (2012). A fast algorithm for matrix balancing. IMA J. Numer. Anal. drs019.

Lieberman-Aiden, E., van Berkum, N.L., Williams, L., Imakaev, M., Ragoczy, T., Telling, A., Amit, I., Lajoie, B.R., Sabo, P.J., Dorschner, M.O., et al. (2009). Comprehensive Mapping of Long-Range Interactions Reveals Folding Principles of the Human Genome. Science 326, 289–293.

